# Genomic, geographic and temporal distributions of SARS-CoV-2 mutations

**DOI:** 10.1101/2020.04.22.055863

**Authors:** Hsin-Chou Yang, Chun-houh Chen, Jen-Hung Wang, Hsiao-Chi Liao, Chih-Ting Yang, Chia-Wei Chen, Yin-Chun Lin, Chiun-How Kao, James C. Liao

## Abstract

The COVID-19 pandemic is the most significant public health issue in recent history. Its causal agent, SARS-CoV-2, has evolved rapidly since its first emergence in December 2019. Mutations in the viral genome have critical impacts on the adaptation of viral strains to the local environment, and may alter the characteristics of viral transmission, disease manifestation, and the efficacy of treatment and vaccination. Using the complete sequences of 1,932 SARS-CoV-2 genomes, we examined the genomic, geographic and temporal distributions of aged, new, and frequent mutations of SARS-CoV-2, and identified six phylogenetic clusters of the strains, which also exhibit a geographic preference in different continents. Mutations in the form of single nucleotide variations (SNVs) provide a direct interpretation for the six phylogenetic clusters. Linkage disequilibrium, haplotype structure, evolutionary process, global distribution of mutations unveiled a sketch of the mutational history. Additionally, we found a positive correlation between the average mutation count and case fatality, and this correlation had strengthened with time, suggesting an important role of SNVs on disease outcomes. This study suggests that SNVs may become an important consideration in virus detection, clinical treatment, drug design, and vaccine development to avoid target shifting, and that continued isolation and sequencing is a crucial component in the fight against this pandemic.

**Significance Statement:** Mutation is the driving force of evolution for viruses like SARS-CoV-2, the causal agent of COVID-19. In this study, we discovered that the genome of SARS-CoV-2 is changing rapidly from the originally isolated form. These mutations have been spreading around the world and caused more than 2.5 million of infected cases and 170 thousands of deaths. We found that fourteen frequent mutations identified in this study can characterize the six main clusters of SARS-CoV-2 strains. In addition, we found the mutation burden is positively correlated with the fatality of COVID-19 patients. Understanding mutations in the SARS-CoV-2 genome will provide useful insight for the design of treatment and vaccination.

## Introduction

SARS-CoV-2 virus has caused the most significant pandemic (COVID-19) in recent history. Within 5 months of its emergence in December 2019, the virus has spread to more than 210 countries, and has caused greater than 170,000 deaths and more than 2.5 million reported cases as of April 22, 2020. This virus is a positive strand RNA virus with a genomic length of about 30K (29,903 nucleotides in the reference genome), and is evolving rapidly. Aged mutations are persisting or diluted away and new mutations are arising. Mutations may increase the fitness of the virus to the environment, while elevating the risk of drug resistance, altering the case fatality rate, and reducing the efficacy of vaccines. Excluding the 5’ leader and 3’ terminal sequences, the genome contains 11 coding regions including S, E, M, N and several open reading frames (ORF1ab, ORF3a, ORF6, ORF7a, ORF7b, ORF8, and ORF10) with various lengths and biological implications. Based on the DNA sequence data of 1,932 SARS-CoV-2 strains from GISAID, NCBI Genbank, and CNCB (data release: March 31, 2020), we performed a phylogenetic analysis based on the full genomes, characterized the geographic and temporal patterns of aged and new mutations, examined genomic profile and identified frequent mutations, inferred linkage disequilibrium (LD) and haplotype structure, constructed the evolutionary paths, correlated phylogenetic clusters with mutations, and investigated how the mutation count influenced the case fatality.

## Results

Based on the whole-genome sequence data, the phylogenetic tree of 1,932 SARS-CoV-2 strains showed there are six major groups (**Figure 1**): (1) Europe-1 (76% of 810 strains are from Europe, with the majority from Iceland); (2) Oceania/Asia (38.6% and 38.6% of 57 strains are from Oceania (all of the Oceania strains are from Australia in this data set) and Asia, respectively, with the majority from Australia and China); (3) Americas (94.4% of 342 strains are from Americas, with the majority from the United States); (4) Europe-2 (80.8% of 104 strains are from Europe, with the majority from Great Britain and Iceland); (5) Asia-1 (72.2% of 133 strains are from Asia, with the majority from China); and (6) Asia-2 (55.5% of 364 strains are from Asia with the majorities from China and Japan).

**Figure 1.**
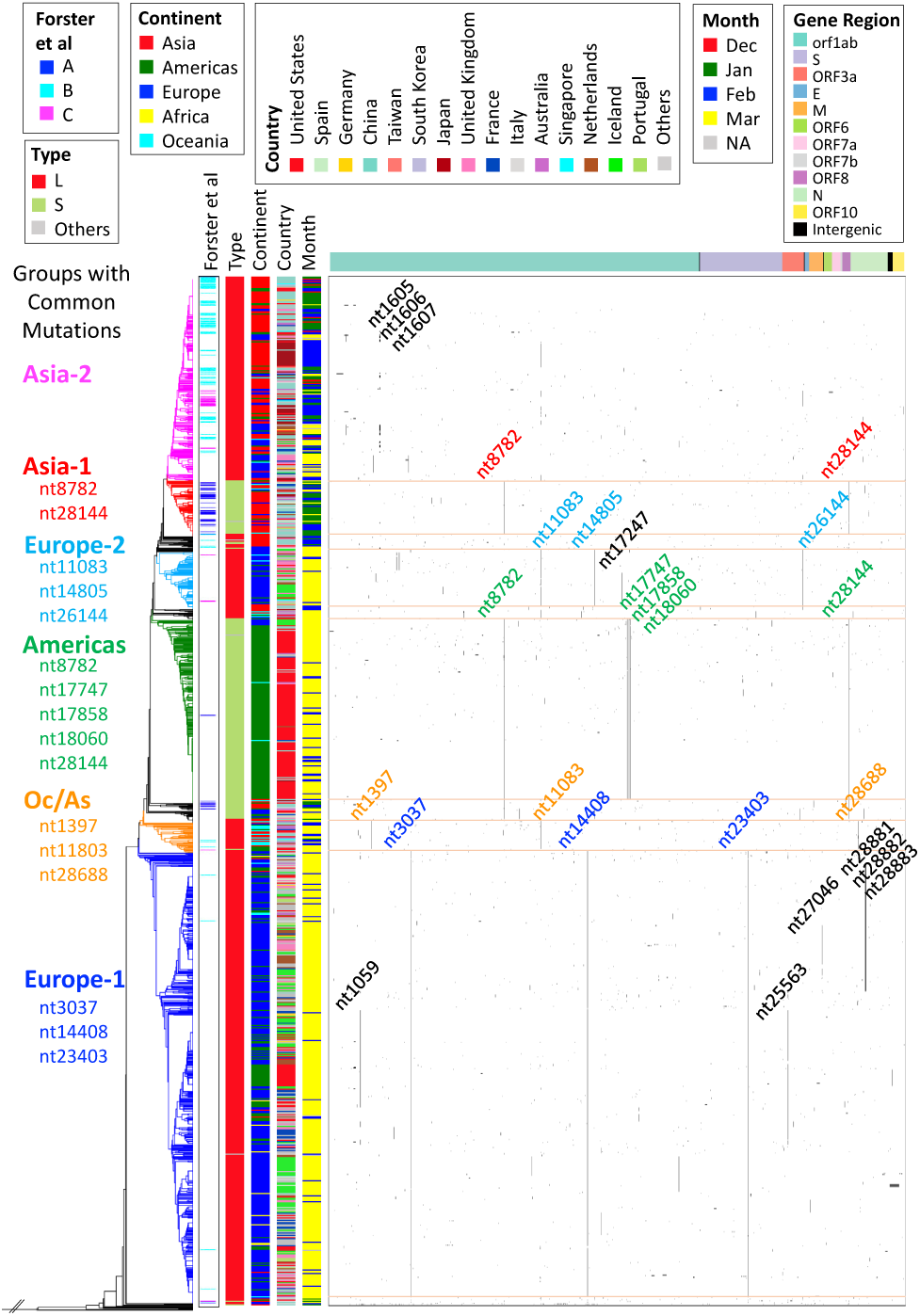
Phylogenetic tree with mutation matrix map of 1,932 SARS-CoV-2 strains. On the right side is the mutation matrix map of 1,932 SARS-CoV-2 strains on 2,139 nucleotides. A black dot at position (*i, j*) indicates an occurrence of a mutation at nucleotide *j* (=1,…, 2139) for virus strain *i* (=1,…, 1932). All 2,139 nucleotides have at least one mutation among 1,932 virus strains. Nucleotides are listed by relative positions in the genome with color bands indicating their corresponding gene regions on the top panel. Five color panels to the left of the mutation matrix were utilized to identify auxiliary information for each virus strain - three strain types (A, B, C) defined by Forster et al. (3), two strain types (L, S) defined by Tang et al (2), and continent, country, with month of data collection. Virus strains were sorted by their relative positions in the phylogenetic tree displayed on the left. Six groups of virus strains were identified via their mutation patterns in the matrix and their relative positions in the phylogenetic tree, Europe-1 (in blue), Oceania/Asia (in orange), Americas (in green), Europe-2 (in cyan), Asia-1 (in red), and Asia-2 (in magenta). Common mutations for each of the six groups of virus strains are also labeled with corresponding colors in the mutation matrix.

We used the strain originally isolated in China (1) as the reference genome and defined variations from the reference as mutations. Among 1,932 SARS-CoV-2 strains studied, the average mutation counts per sample in Europe and Americas were much higher than that in Asia (**Figure 2A**). This phenomenon partially reflects the early occurrence of epidemic of COVID-19 in Asia. Among the nations having at least 10 cases reported in this dataset, the top three nations with the highest average mutation counts were all located in Europe: Spain, Belgium, and Finland. The three nations with the lowest value were located in Asia: Singapore, Japan, and China. The six major clusters derived from the phylogenetic analysis (**Figure 1**) also showed more mutations in the European and American clusters compared to the two Asian clusters. The average mutation counts per sample in Europe-1, Oceania/Asia, Americas, Europe-2, Asia-1, and Asia-2 were 6.60, 6.46, 6.67, 6.21, 4.07, and 2.68, respectively.

**Figure 2A.**
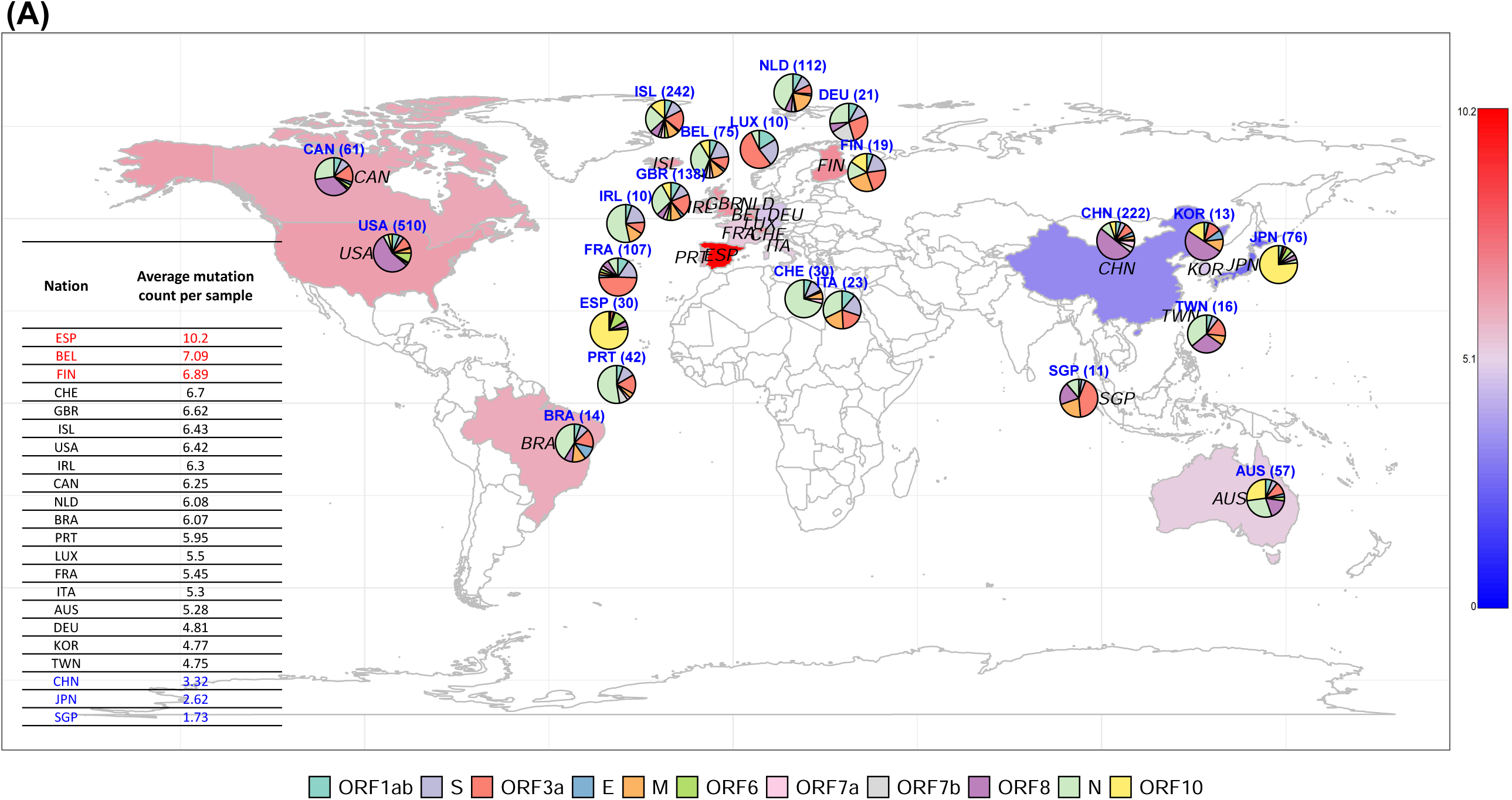
WGeographic distributions of the average mutation counts in the entire genome and the standardized mutation densities in the eleven gene regions globally. A mutation is defined by a nucleotide change from the original nucleotide in the reference genome to the alternative nucleotide in the studied viral genomes. In each nation, the average mutation count per sample (i.e., the number of mutations in the genomes of all virus strains divided by the number of strains) is displayed in a color spectrum from blue (low average mutation count) to red (high average mutation count). The statistics of the average mutation count per sample are provided. Mutation density in each of the eleven gene regions (i.e., the number of mutations divided by the number of nucleotides in a gene region for all virus strains) is standardized (i.e., the mutation density in each gene divided by the sum of mutation densities in the eleven gene regions) and shown in a pie chart. A table of the average mutation count per sample for the nations having at least 10 cases reported in this dataset is listed.

Mutations were distributed across the genome in different patterns according to the geographic locations that the samples were isolated. Mutation densities (i.e., the number of mutations per nucleotides in the gene region) were high in several gene regions including ORF3a (Luxembourg, France, and Singapore), ORF8 (the United States, Korea, and China), N (Switzerland, Ireland, and Portugal), and ORF10 (Japan and Spain) (**Figure 2A**). ORF1ab harbors a significant number of mutations, but because of its length the mutation density appears to be low. The United States had the largest sample size (N = 510) and the average mutation count was 6.42. Minnesota and Arizona showed the highest and lowest average mutation counts in the US (**Figure S1**), respectively. The dominant mutation types in the east (NY) and the west (CA, WA, and UT) were different, with mutations in ORF3a dominating in the former, while mutations in ORF8 in the latter states.

We found a total of 169,060 mutation counts. The average mutation count per locus was 169,060/29,903 = 5.654. The average mutation count per sample was 169,060/1,931 = 87.55. The average mutation count per locus per sample was 0.298%. After removing the mutations in 5’ leader and 3’ terminal sequences, the total mutation counts for all samples across genome reduced to 11,223. The average mutation count per locus was 11,223/29,409 = 0.3816. The average mutation count per sample was 11,223/1931 = 5.812. The average mutation count per locus per sample was 0.0198%.

Globally, the average mutation count per sample was increasing with time (r = 0.65 in a linear regression) (**Figure 2B**). This result indicates that the mutated strains persisted and were expanding. A large number of the mutations originated from China at the early stage have overwhelmed the samples collected from Americas and Europe after a few months. In addition to the cumulated aged mutations, we also identified a significant number of new mutations occurring at each date. Unlike the increasing trend of the cumulated aged mutations, the number of new mutations occurring globally at each date remained at a constant level. The major contributor of the new mutations was China at the early stage but shifted to Americas and Europe after mid-February (**Figure 2C**).

**Figure 2B.**
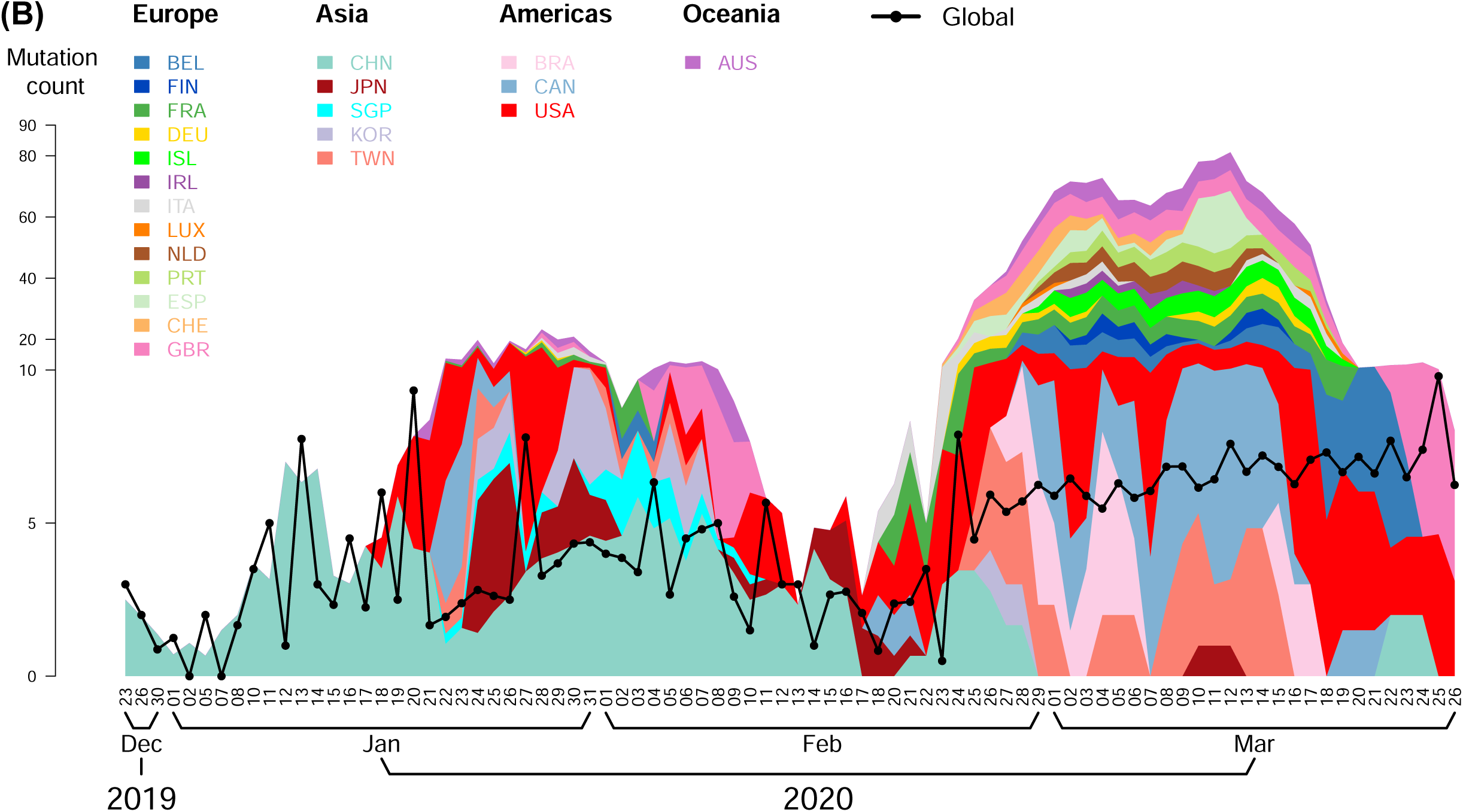
Temporal trend of the average mutation counts in globe and nations. In each nation, the average mutation count per sample (i.e., the number of mutations in the genomes of all virus strains divided by the number of strains) was calculated. The black dotted line indicates the trajectory of the average mutation counts in globe. A fitted regression line is drawn.

**Figure 2C.**
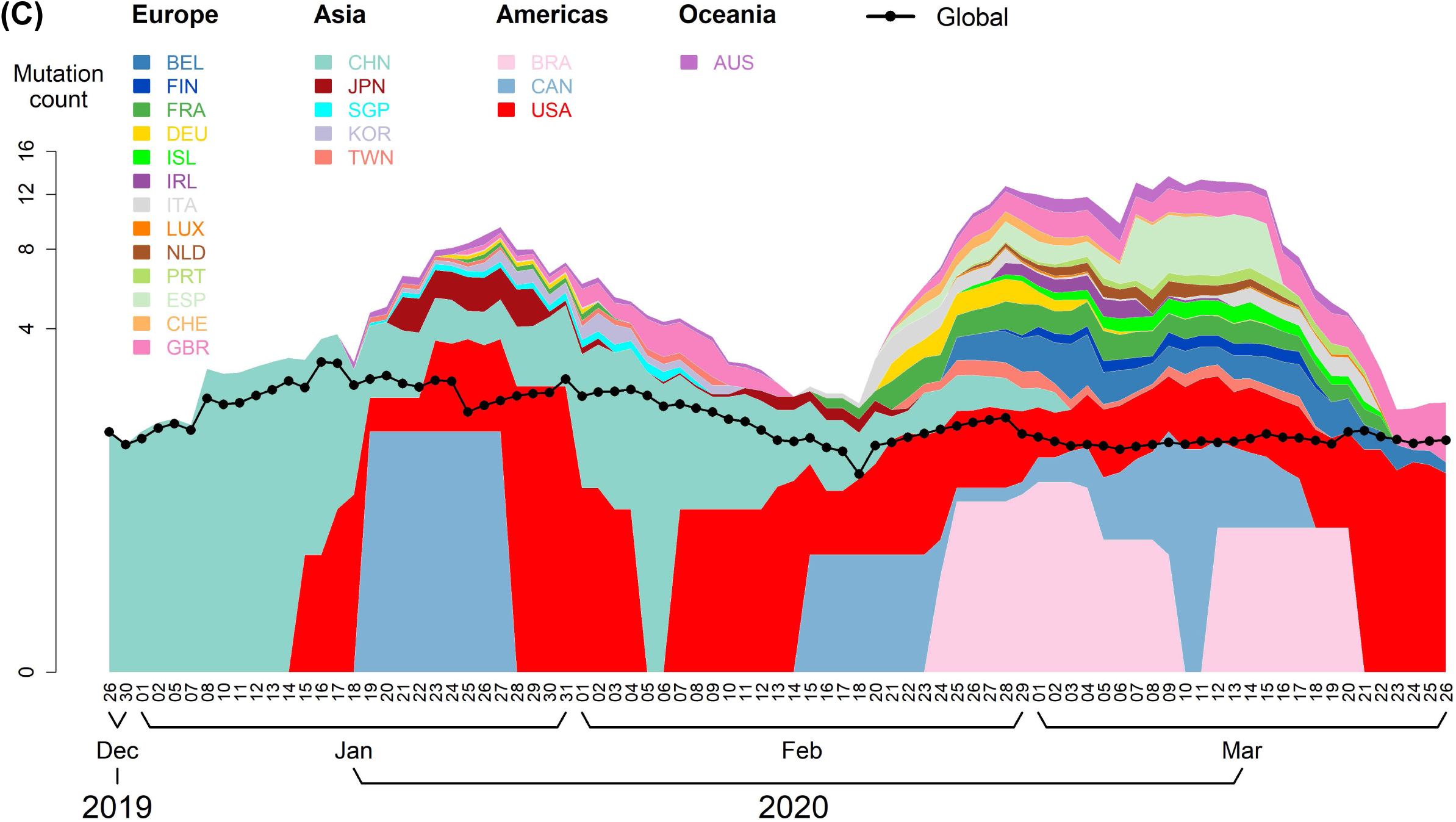
Temporal trend of the new mutations in globe and nations. (A) A new mutation is defined by a nucleotide change from the original nucleotide in the reference genome to the alternative nucleotide in the studied viral genomes and had never been observed before the specific time point in this dataset. The black dotted line indicates the trajectory of the average counts of new mutations in globe. A fitted regression line is drawn.

**Figure 2D.**
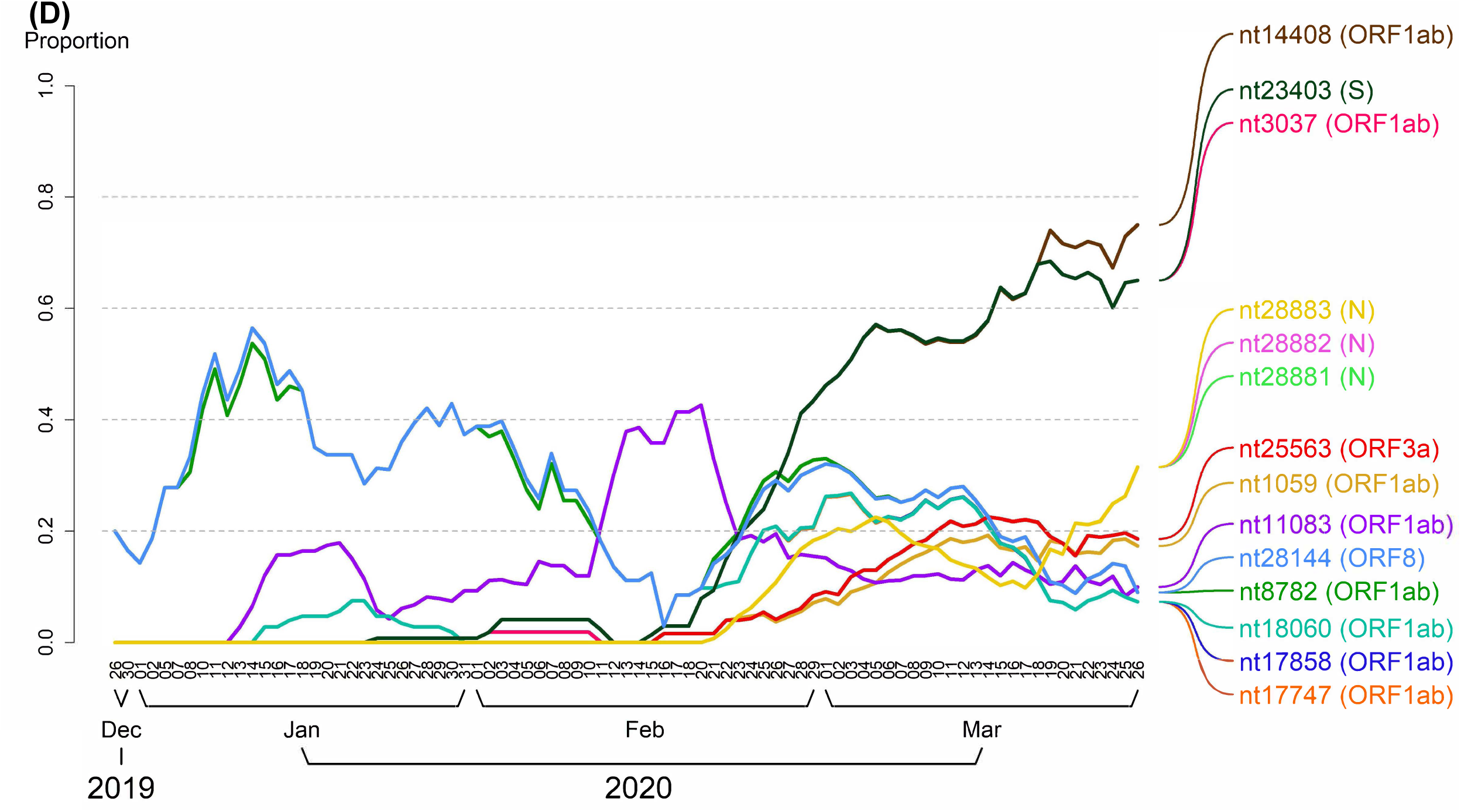
Temporal frequency profile of the frequent mutations. In each time point of data collection, the average mutation count per sample (i.e., the number of mutations occurred at the nucleotide in all virus strains divided by the number of strains) is displayed. Only the fourteen frequent mutations with a frequency of >0.1 is displayed.

Fourteen frequent mutations with a mutation frequency of >0.1 were identified (**Table S1** and **Figure 3A**). These frequent mutations showed interesting patterns. First, the mutations have a similar mutation rate if they were first observed at the same or close dates and in strong LD. For example, nt8782 (C to T) and nt28144 (T to C) that were used to define L type and S type of SARS-CoV-2 (2) first co-appeared on Jan 5th 2020 and the coefficient of LD is R2 = 0.983 (**Figure 3A**). The reference strain used here is the L type, while the S type exhibits both mutations. This implies that the frequent mutations on the same haplotype were co-transmitted to the infected cases during infection. The haplotypes were located in the same or across different gene regions. Other cases first co-observed at the same date included nt1059 (C to T) and nt25563 (G to T) (Feb 21st 2020); nt28881 (G to A), nt28882 (G to A), and nt28883 (G to C) (Feb 25th 2020). Nucleotides nt3037 (C to T), nt14408 (C to T), and nt23403 (A to G) having high pairwise LD are the most frequent nucleotides in the current data set. Nucleotides nt17747 (C to T), nt17858 (A to G), and nt18060 (C to T) having almost identical nucleotide frequency also showed high pairwise LD. Nucleotide nt11083 had two alternative nucleotides (G to T or C) and was not correlated to the other frequent mutations.

**Figure 3A.**
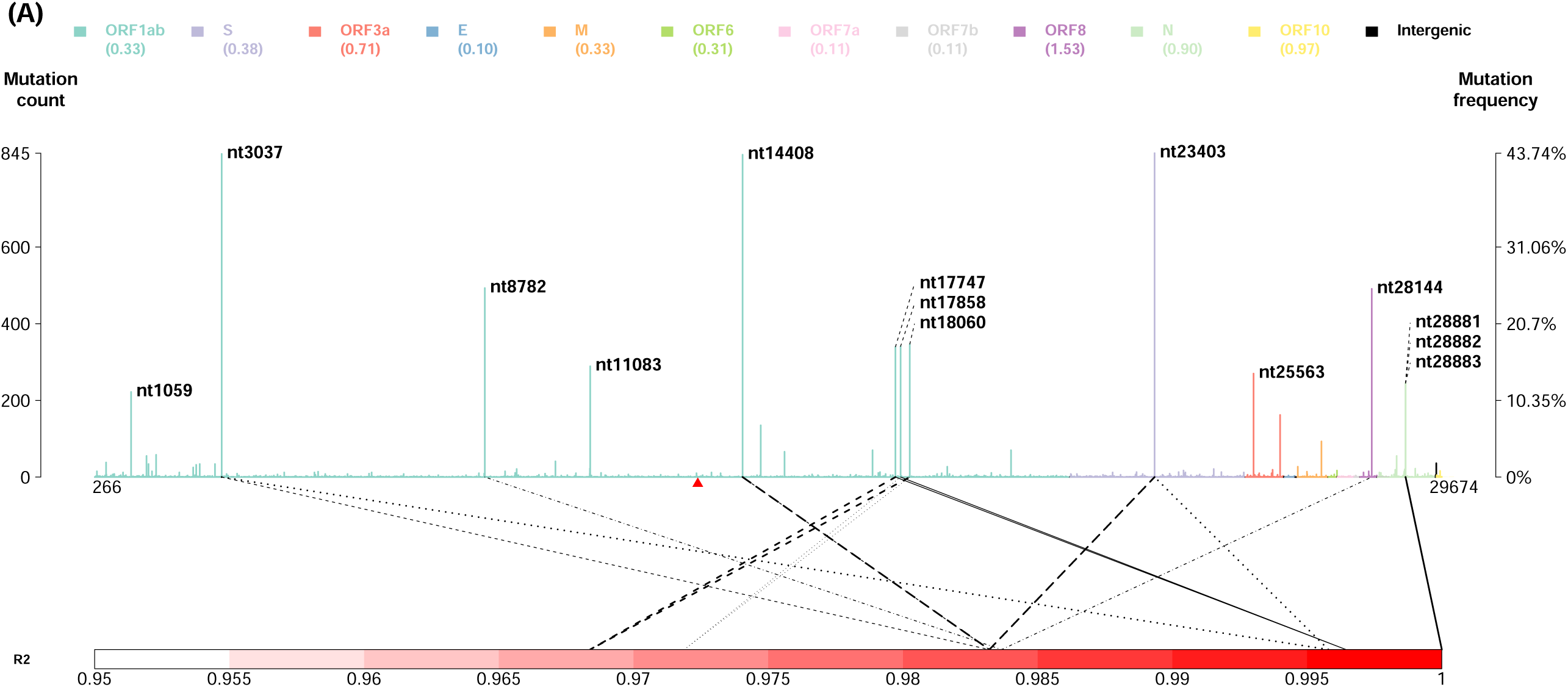
Genomic profile of the average mutation counts across the viral genome. In each nucleotide, the average mutation count per sample (i.e., the number of mutations occurred at the nucleotide in all virus strains divided by the number of strains) is displayed. Nucleotides in different gene regions are displayed in different color. A triangle mark indicates the starting site of −1 ribosomal frameshift signal in ORF1ab. Two ends (5’ leader and 3’ terminal sequences) are not shown. Pairwise linkage disequilibrium (R2) of the frequent mutations is shown. Only the pairs of frequent mutations with a frequency of >0.1 and an R2 value of >0.95 are shown. The same line type is used to indicate a pair of mutations.

Second, the evolutionary path of the fourteen frequent mutations was constructed (**Figure 3B** and **3C**) based on their first observation time. Note that the real occurrence date of mutations should be earlier because of left censoring of mutation events. The haplotype frequency of the reference strain without any of the fourteen frequent mutations was 16.86% and was exceeded by several mutated haplotypes (**Figure 3C**). The haplotype frequencies varied across continents and nations. For example, the mutation haplotype 8782T-28144C was prevalent in the US but mutation 11083C/T and mutation haplotype 28881A-28882A-28883C were not), reflecting the different transmission paths of viral strains and their different ability to adapt to the local environment (**Figure 3D**).

**Figure 3B.**
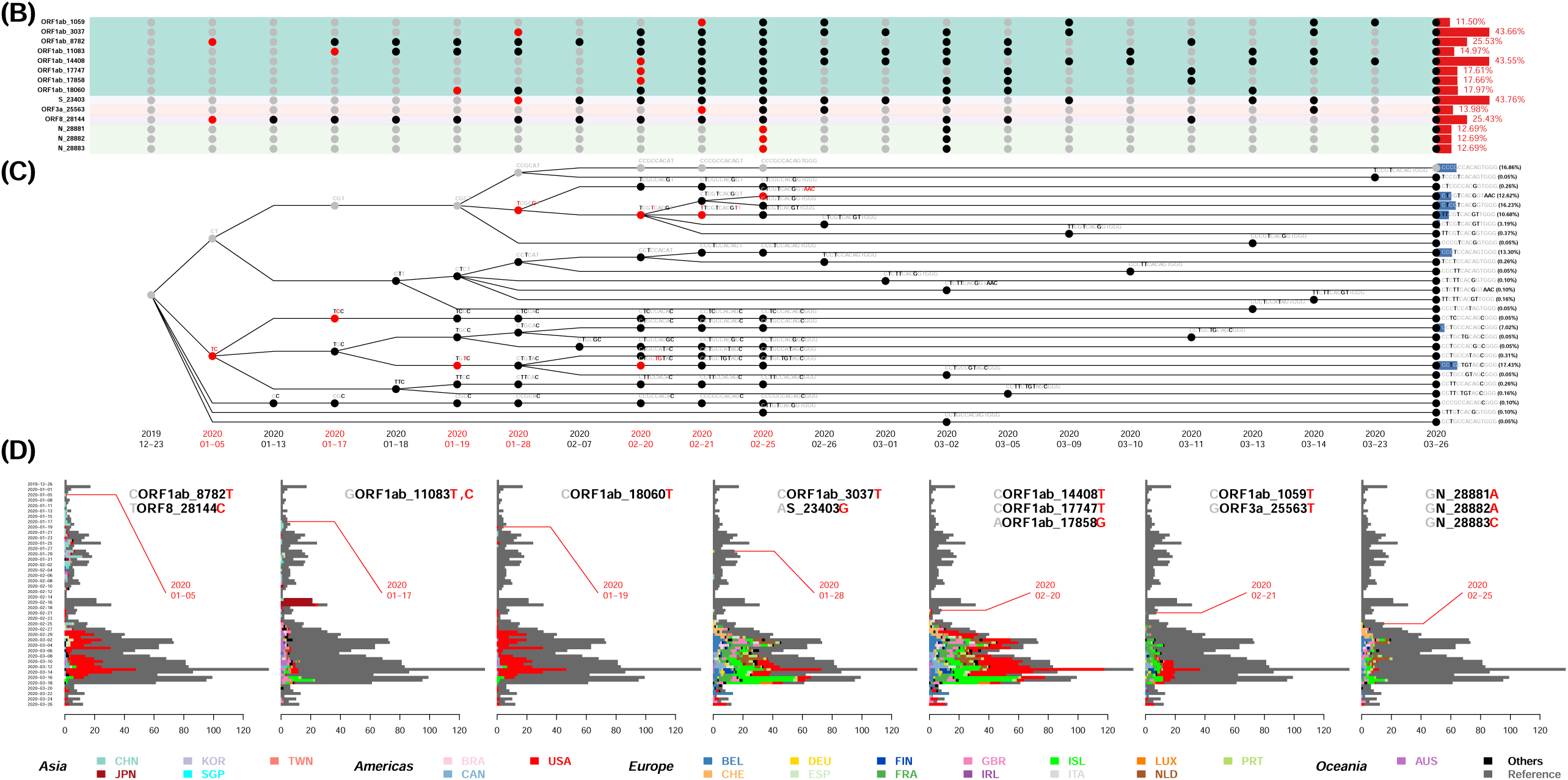
Evolutionary path of the prevalent mutations. Evolutionary path of the fourteen prevalent mutations with a frequency of >0.1 are displayed. The color scheme used to indicate gene regions is the same as that in Figure 1. A red dot indicates the first time to observe the mutation. A black dot indicates the following mutation(s) of the mutation. A gray dot indicates the nucleotide is the same as the nucleotide in the reference genome. The average mutation count per sample in this data set is shown in a histogram in the right-hand side of this subfigure. **Figure 3C. Haplotype tree of the prevalent mutations**. Haplotype tree of the fourteen prevalent mutations with a frequency of >0.1 are displayed. The average count of mutation haplotype per sample in this data set are shown in a histogram in the right-hand side of this subfigure. **Figure 3D. Geographic temporal frequency distribution of the prevalent mutations**. Geographic and temporal frequency distributions of the fourteen prevalent mutations with a frequency of >0.1 are displayed. For each frequent mutation, the average mutation counts by the data collection time are shown in a stacked bar chart that the frequencies in different nations are indicated by different colors.

It is also interesting to see the time trajectory of the average mutation count in the fourteen frequent mutations (**Figure 2D**). Nucleotides nt14408, nt23403 and nt3037 were in strong LD and have emerged to be the most prevalent mutations. The raising patterns of the average mutation count may imply natural selection and adaptation to the different environment. Unexpectedly, the earliest mutations reported were in nt8782 and nt28144 in this data set, also with strong LD, but their frequency has gradually declined. Strains with the relatively new mutations in nt14408, nt23403 and nt3037 have emerged to become the most common strains identified in the world, particularly in subgroup Europe-1.

The major clusters of SARS-CoV-2 strains identified in the phylogenetic tree can be characterized by the key mutations (**Figure 1**), most of which were also shown in **Figure 2D**. Two European groups had quite different mutation patterns. The Europe-1 group carried the specific mutations nt3037T (ORF1ab), nt14408T (ORF1ab), and nt23403G (S), in addition to the nt241T mutation in 5’ leader sequence. These mutations were hardly observed in other groups. The Europe-2 group is characterized by mutations nt11083T (ORF1ab), nt14805T (ORF1ab), and nt26144T (ORF3a). The Oceania/Asia group carried mutations nt1397A (ORF1ab), nt11083T (ORF1ab) and nt28688C (N). The Americas group carried mutations nt8782T (ORF1ab) and nt28144C (ORF8), which were used to define the S sub-type (2). These two mutations were also observed in the Asia-1 group, but not in the other groups. Additionally, the Americas group also carried mutations nt17747T (ORF1ab), nt17858G (ORF1ab), and nt18060T (ORF1ab). The Asia-2 group is distinct from others because the strains in this group did not carry most of the frequent mutations. On average, the dates of sample collection of the two Asian groups were closest to the first emergence of COVID-19 in December 2019 and their genomes were closest to the reference. They were characterized by multiple mutations with low frequencies. Only about 6% of viral strains cannot be categorized into the six main phylogenetic clusters.

Finally, a point worth noting is the strong linear correlation between the case fatality rate and the average mutation count per sample (r = 0.4258, p = 0.0482) as of April 9, 2020 (**Figure 4A**). Among eleven gene regions, only ORF1ab showed a significant linear correlation (r = 0.407, p = 0.0542) (**Figure 4B**), suggesting that ORF1ab mutation may contribute the most to the case fatality rate. Surprisingly, by April 21, 2020 the correlation between case fatality and the average mutation count has become even more significant with p= 0.0348 (**Figure 4C**), and its contribution from ORF1ab has a p value of 0.0291 (**Figure 4D**). These results indicate that mutation has already impacted clinical outcome of COVID-19, not just a viral fitness evolution event.

**Figure 4.**
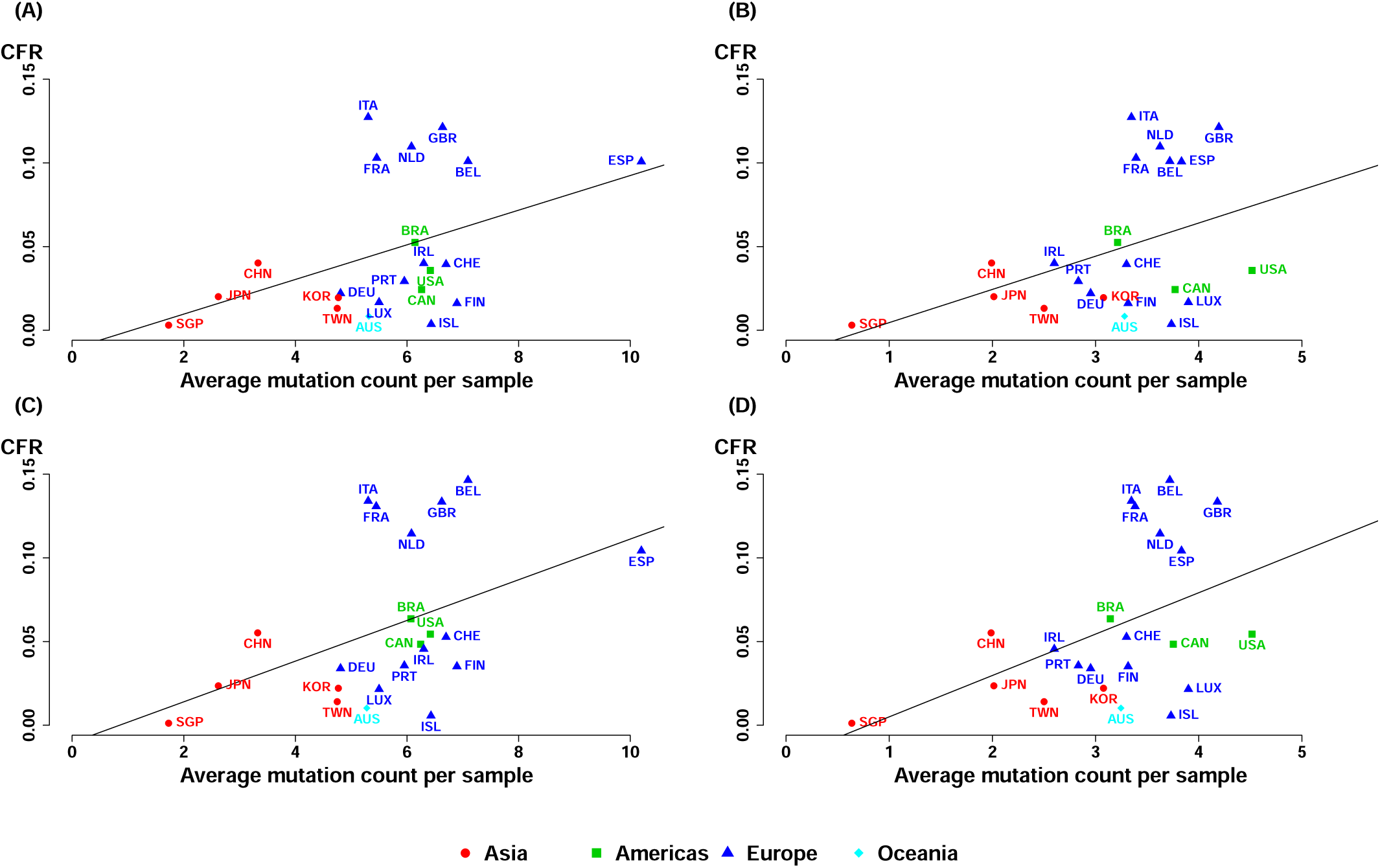
Relationship of the case fatality rate and mutation burden. (A) Vertical axis indicates the case fatality rate and horizontal axis indicates the mutation burden (the average mutation counts per sample in the nation). Each point indicates a nation. Different colored symbols indicate continents. A linear regression line is drawn. (A) Mutations in the entire genome (date: April 9, 2020); (B) Mutations in ORF1ab (date: April 9, 2020); (C) Mutations in the entire genome (date: April 21, 2020); (D) Mutations in ORF1ab (date: April 21, 2020).

## Discussion

In summary, we report evidence for time and geographic variations of SARS-CoV-2 mutations, and identified six major subgroups of SARS-CoV-2 strains with strong geographic preferences based on the complete genomes of 1,932 SARS-CoV-2 strains. These subgroups can be characterized by 14 common mutations, most of which occurred in ORF1ab, with notable exceptions in S, N, ORF3a and ORF8. This result suggests the importance of these genes in viral fitness and perhaps clinical relevance. Unexpectedly, we found that the case fatality rate in each country correlates positively with the average mutation count per sample. The positive correlation was contributed mainly from gene region ORF1ab, which codes for polyproteins that are cleaved to become proteases, RNA-dependent RNA polymerase, helicase, and several non-structural proteins (nsp). The significance of the correlation increased with time, suggesting that mutation may have impacted viral transmission, clinical manifestation, or treatment outcomes.

Compared to the L and S types originally reported (2), and the A, B, C types reported recently (3), our classification of six subgroups provides a mutation-based taxonomy of viral strains and explains the heterogeneity of strains within each of L and S types and A, B, C types. Interestingly, 5 out of 6 subgroups are characterized by a few common mutations in each subgroup. This concept and method for classification and characterization of viral strains can be applied to other viruses of public health concerns. As more whole genome sequencing data of SARS-CoV-2 become available online, we will be better-posed to decipher and understand genomic, geographic and temporal distributions of viral mutations. However, representativeness of the submitted genomic data should be considered carefully when explaining the results.

The different protein reading frames exhibited different mutation frequencies depending on the geographic location. We dissected the prevalent mutations, investigated their LD structure, traced their evolutionary path, and found their importance in the characterization of phylogenetic clusters. Interestingly, many frequent mutations occurred simultaneously, with LD close to 1. These mutations may occur simultaneously because of strong positive interactions. Alternatively, they might occur sequentially but appeared simultaneously due to insufficient sampling. The biological implication of each mutation and their interactions remain an interesting topic to be explored. As aged mutations continued to accumulate in time and expand globally, new mutations emerged in the gene pool constantly to enrich the diversity of mutations. Since the fatality rate appears to correlate with the mutation count, the continued isolation and sequencing of the viral genome to monitor mutations become a crucially important component in the fight against this pandemic.

## Materials and Methods

We downloaded the whole-genome sequence data from the Global Initiative on Sharing Avian Influenza Data (GISAID) database (https://www.gisaid.org/), National Center for Biotechnology Information (NCBI) Genbank (https://www.ncbi.nlm.nih.gov/genbank/), and China National Center for Bioinformation (CNCB) (https://bigd.big.ac.cn/ncov/release_genome) on Mar 31st 2020. After discarding the replicated sequences in the three databases and the sequences with a low quality indicator or no quality information, it remained the complete sequences of 1,938 SARS-CoV-2 genomes, including the Wuhan-Hu-1 reference genome with 29,903 nucleotides. Multiple sequence alignment was performed by using MUSCLE (4). Mutation was identified as variation from the reference. Generalized association plot (GAP) (5) was used to visualize the mutation patterns and identify the outliers in variations and samples (**Figure S2**). We removed two ends (5’ leader and 3’ terminal sequences) of the SARS-CoV-2 genome because of a significant number of gaps. That is, we focused on nucleotides from positions 266 to 29,674. Before removing the two ends, we observed two nucleotides with a non-negligible mutation frequency relative to their neighboring regions: nt241 with a mutation frequency of 45.5% (and this nucleotide was in high LD with nt3037, nt14408, and nt23403) and nt29742 with a mutation frequency of 5.3%. We also removed four samples with a large deletion in ORF8 (sample EPI_ISL_417518 from Taiwan with EPI_ISL_141378, EPI_ISL_141379, and EPI_ISL_141380 from Singapore), sample EPI_ISL_415435 from UK with a large deletion in ORF1ab, and sample EPI_ISL_413752 from China with a large number of deletions (>300 nucleotides).

A new mutation at time *t* was defined as a mutation that has never been observed before *t* in this data set. Average mutation count per sample and/or per locus were calculated for nations and for the data collection time points to study geographic and temporal distributions of mutations. Mutation frequencies in gene regions were illustrated and coefficient of LD (R2) between pairs of nucleotides was calculated by using PLINK (6). Frequent mutations were defined as mutations with a frequency of >0.1 in this paper. Annotation of the frequent mutations was collected from China National Center for Bioinformation. Nucleotide frequency and haplotype frequency were calculated by using a direct counting method. Phylogenetic tree analysis was performed by using MEGA X (7). GAP was applied to present the relationship between phylogenetic cluster and mutations. Finally, the data of case fatality rate was collected the website of Coronavirus Resource Center, Johns Hopkins University on Apr 9th 2020 and Apr 21st 2020. A linear regression model was built to correlate the average mutation count per sample and case fatality rate. Other statistical graphs were generated using our self-developed R programs.

## Supplemental Information

**Table S1.** Annotation of the fourteen frequent mutations.

| Nucleotide (Gene) | Base change | Annotation type | Protein:Position | Amino acids change | Codon number:<br>Codon change | Alternate allele frequency |
| --- | --- | --- | --- | --- | --- | --- |
| nt1059 (ORF1ab) | C -> T | missense variant | QHD43415.1:p.265 | T>I | c.794:aCc>aTc | 0.11 |
| nt3037 (ORF1ab) | C -> T | synonymous variant | QHD43415.1:p.924 | F | c.2772:ttC>ttT | 0.43 |
| nt8782 (ORF1ab) | C -> T | synonymous variant | QHD43415.1:p.2839 | S | c.8517:agC>agT | 0.25 |
| nt11083 (ORF1ab) | G -> T;<br>G -> C | missense variant;<br>coding sequence variant | QHD43415.1:p.3606 | L>F;<br>- | c.10818:ttG>ttT;<br>c.10818:ttG>ttC | 0.14;<br>0.00 |
| nt14408 (ORF1ab) | C -> T | missense variant | QHD43415.1:p.4715 | P>L | c.14144:cCt>cTt | 0.43 |
| nt17747 (ORF1ab) | C -> T | missense variant | QHD43415.1:p.5828 | P>L | c.17483:cCt>cTt | 0.18 |
| nt17858 (ORF1ab) | A -> G | missense variant | QHD43415.1:p.5865 | Y>C | c.17594:tAt>tGt | 0.18 |
| nt18060 (ORF1ab) | C -> T | synonymous variant | QHD43415.1:p.5932 | L | c.17796:ctC>ctT | 0.18 |
| nt23403 (S) | A -> G | missense variant | QHD43416.1:p.614 | D>G | c.1841:gAt>gGt | 0.44 |
| nt25563 (ORF3a) | G -> T | missense variant | QHD43417.1:p.57 | Q>H | c.171:caG>caT | 0.14 |
| nt28144 (ORF8) | T -> C | missense variant | QHD43422.1:p.84 | L>S | c.251:tTa>tCa | 0.25 |
| nt28881 (N) | G -> A | missense variant | QHD43423.2:p.203 | R>K | c.608:aGg>aAg | 0.13 |
| nt28882 (N) | G -> A | synonymous variant | QHD43423.2:p.203 | R | c.609:agG>agA | 0.13 |
| nt28883 (N) | G -> C | missense variant | QHD43423.2:p.204 | G>R | c.610:Gga>Cga | 0.13 |

**Figure S1.**
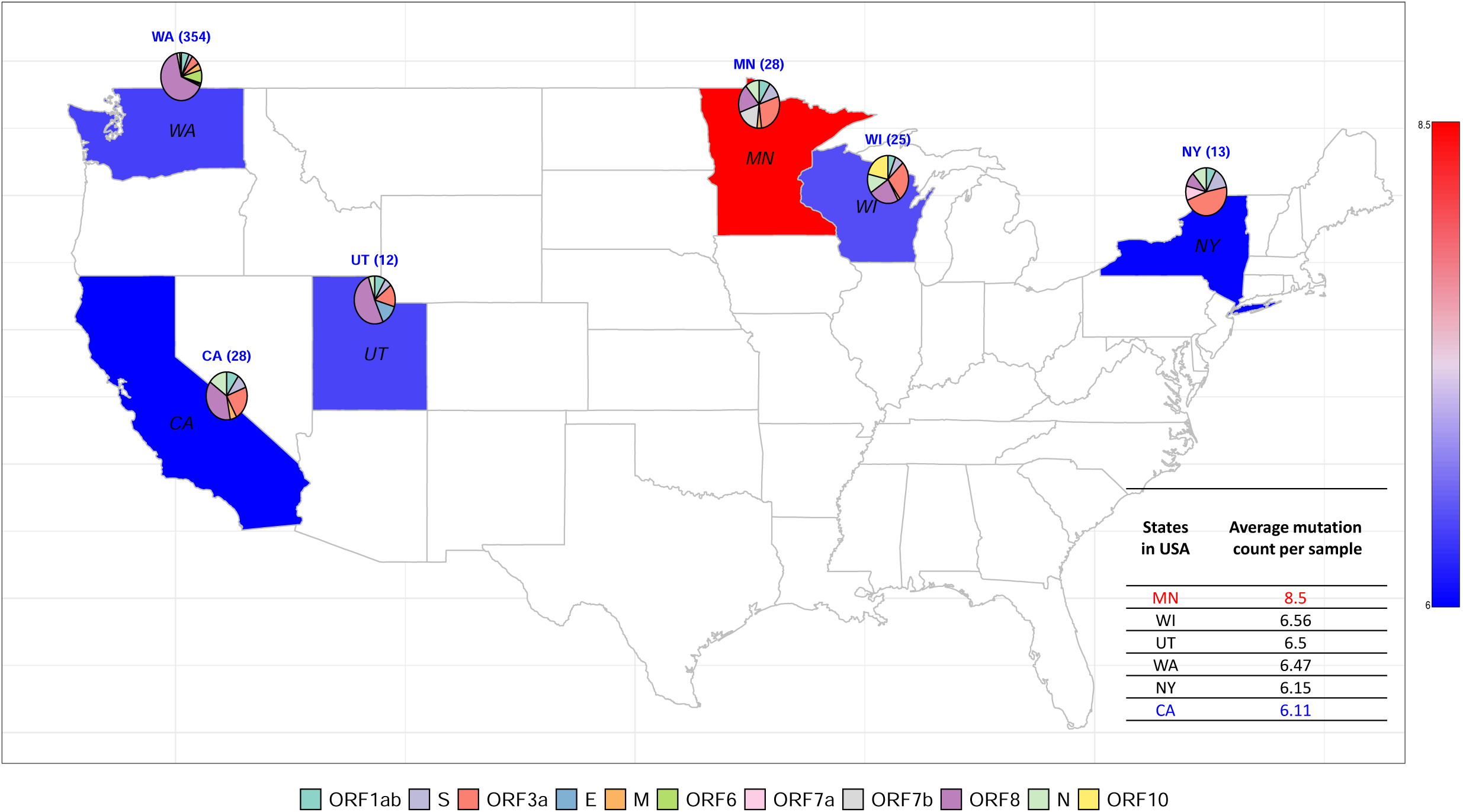
Geographic distributions of the average mutation counts in the entire genome and the standardized mutation densities in the eleven gene regions in the United States. A mutation is defined by a nucleotide change from the original nucleotide in the reference genome to the alternative nucleotide in the studied viral genomes. In each nation, the average mutation count per sample (i.e., the number of mutations in the genomes of all virus strains divided by the number of strains) is displayed in a color spectrum from blue (low average mutation count) to red (high average mutation count). The statistics of the average mutation count per sample are provided. Mutation density in each of the eleven gene regions (i.e., the number of mutations divided by the number of nucleotides in a gene region for all virus strains) is standardized (i.e., the mutation density in each gene divided by the sum of mutation densities in the eleven gene regions) and shown in a pie chart. A table of the average mutation count per sample for the states having at least 10 cases reported in this dataset is listed.

**Figure S2.**
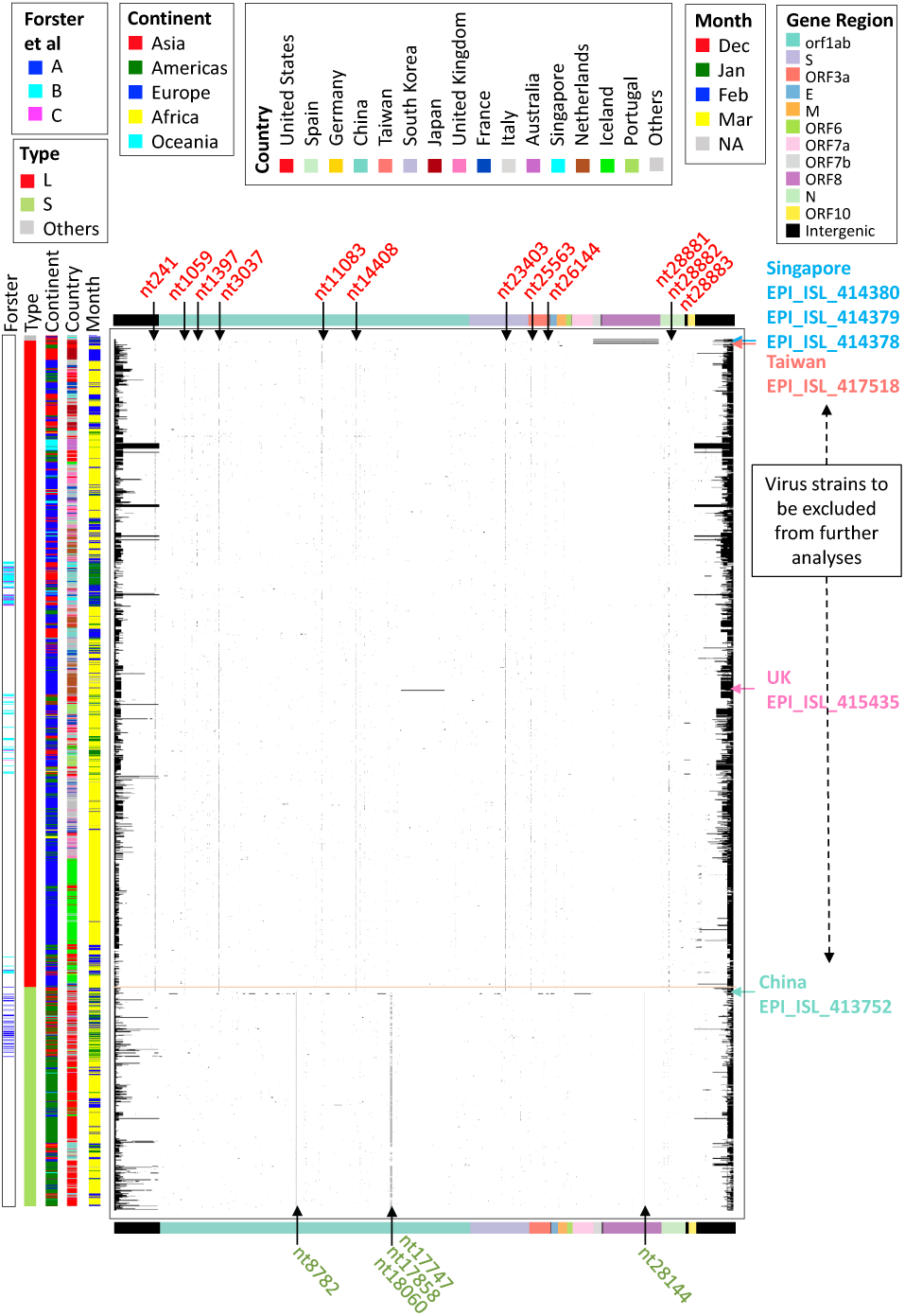
Mutation matrix map of 1,938 SARS-CoV-2 strains on 3,607 nucleotides. A black dot at position (*i, j*) indicates an occurrence of a mutation at nucleotide *j* (=1,…, 3607) for virus strain *i* (1,…, 1938). All 3,607 nucleotides have at least one mutation among 1,938 virus strains. Nucleotides are listed by relative positions in the genome with color bands indicating their corresponding gene regions on the top and bottom panels. Five color panels on the left were utilized to identify auxiliary information for each virus strain - three strain types (A, B, C) defined by Forster et al. (3), two strain types (L, S) defined by Tang et al (2), and continent, country, with month of data collection. Virus strains were sorted by the two types defined in Tang et al, type L (in red), type S (in light green). There is a small group of non-S/L type strains (in grey) listed at the first few rows of the matrix. Nucleotides labeled in red (green) on top (at bottom) of the matrix were type L (S) specific mutations. Listed on the right of the matrix are six virus strains that exhibit unusual mutation patterns and will be excluded from further analyses.

## References

1. F. Wu et al., A new coronavirus associated with human respiratory disease in China. Nature 579, 265–269 (2020).

2. X. Tang et al., On the origin and continuing evolution of SARS-CoV-2. National Science Review (2020).

3. P. Forster, L. Forster, C. Renfrew, M. Forster, Phylogenetic network analysis of SARS-CoV-2 genomes. Proc Natl Acad Sci U S A 10.1073/pnas.2004999117 (2020).

4. R. C. Edgar, MUSCLE: multiple sequence alignment with high accuracy and high throughput. Nucleic Acids Res 32, 1792–1797 (2004).

5. H. M. Wu, Y. J. Tien, C. H. Chen, GAP: A graphical environment for matrix visualization and cluster analysis. Comput. Stat. Data Anal. 54, 767–778 (2010).

6. S. Purcell et al., PLINK: a tool set for whole-genome association and population-based linkage analyses. Am. J. Hum. Genet. 81, 559–575 (2007).

7. S. Kumar, G. Stecher, M. Li, C. Knyaz, K. Tamura, MEGA X: Molecular Evolutionary Genetics Analysis across Computing Platforms. Mol Biol Evol 35, 1547–1549 (2018).

